# Brief encounters with real objects modulate medial parietal but not occipitotemporal cortex

**DOI:** 10.1101/2024.08.05.606667

**Authors:** Susan G. Wardle, Beth Rispoli, Vinai Roopchansingh, Chris I. Baker

## Abstract

Humans are skilled at recognizing everyday objects from pictures, even if we have never encountered the depicted object in real life. But if we have encountered an object, how does that real-world experience affect the representation of its photographic image in the human brain? We developed a paradigm that involved brief real-world manual exploration of everyday objects prior to the measurement of brain activity with fMRI while viewing pictures of those objects. We found that while object-responsive regions in lateral occipital and ventral temporal cortex were visually driven and contained highly invariant representations of specific objects, those representations were not modulated by this brief real-world exploration. However, there was an effect of visual experience in object-responsive regions in the form of repetition suppression of the BOLD response over repeated presentations of the object images. Real-world experience with an object did, however, produce foci of increased activation in medial parietal and posterior cingulate cortex, regions that have previously been associated with the encoding and retrieval of remembered items in explicit memory paradigms. Our discovery that these regions are engaged during spontaneous recognition of real-world objects from their 2D image demonstrates that modulation of activity in medial regions by familiarity is neither stimulus nor task-specific. Overall, our results support separable coding in the human brain of the visual appearance of an object from the associations gained via real-world experience. The richness of object representations beyond their photographic image has important implications for understanding object recognition in both the human brain and in computational models.

## Main

Humans excel at visual object recognition — we can recognize everyday objects in photographs or from simplified line drawings rapidly and with ease^1,2^. Deep neural networks trained on object recognition are also able to label objects with remarkable accuracy, now rivaling human performance^3,4^. However, when humans look at a photograph, we are capable of much more than simply labelling the objects in the image, e.g. as a “dog”. We can infer other things about the depicted object based on experience (“Labradors are friendly”) or visual features of the image (“the dog is wet”). If we have personal experience with the photographed object (such as a photo of our own dog), we also have access to another form of object knowledge based on real-world experience (e.g. memory of a particular walk, knowledge about the dog’s preference for peanut butter). Much of the everyday human experience involves physically interacting with familiar three-dimensional objects, yet most empirical studies of human object recognition have relied on two-dimensional images of unfamiliar objects^5^. Here we investigate to what extent real-world experience with an object influences the neural representation of its two-dimensional image in the human brain.

Given that a major goal in cognitive neuroscience is to understand real-world object recognition outside of the laboratory and beyond the computer screen^5^, revealing how real-world experience with an object shapes the representation of its image in the brain is important for contextualizing the large body of existing research on the neural representation of object images^6–10^. Previous research using two-dimensional images of objects has identified regions in lateral occipital (LO) and ventral temporal cortex (VTC) that respond preferentially to images of intact objects compared to scrambled versions of the same images^8,11,12^. These regions are sensitive to object shape in photographs^12^, but also respond to more schematic depictions of objects such as line drawings^13^ and shape from texture and motion cues^14^. Prior research with real 3D objects has shown that they are processed differently than images of objects^5^, both behaviorally^15–18^ and in the human brain^19,20^. For example, repetition suppression in object-responsive cortex for repeated presentations of real objects is weaker than for object images^20^.

Although we can extract a substantial amount of information from two-dimensional images of objects, real world experience with a specific object in three dimensions is likely to support multifaceted object representations that are not tied to their two-dimensional image. For example, for a given object, the projected retinal image is vastly different across encounters because of variability in visual characteristics such as lighting, position, proximity and viewing angle. Recognizing an object across these substantial visual transformations involves a complex computation to tease apart the stable visual features of the object from the changeable visual properties of the viewing context, and a large amount of research with two-dimensional images has been directed at understanding how this classic “invariance problem” of object recognition is solved^20^. In our everyday interactions with real-world objects, we view them from multiple angles and under different viewing conditions. Thus one way that real-world experience with an object may manifest in object-responsive visual cortex is in more invariant object representations.

To determine whether real-world experience with an object influences the neural representation of its photographic image, we developed an experimental paradigm that combined real-world object exploration with functional magnetic resonance imaging (fMRI) and machine learning analysis methods (**Fig. 1a**). Participants freely explored everyday objects such as mugs and toys prior to having their brain activity recorded with fMRI while looking at photographs of those same objects, as well as a matched set of unexplored objects of the same type. For an ecologically valid test of object invariance, we included two different photographs taken of each object in different contexts (indoor vs. outdoor). Although the invariance of object representations in ventral occipitotemporal cortex has frequently been studied^22,23^, it still unknown whether different photographs of the same object are represented similarly in these regions. This is because studies have typically focused on the level of object categories rather than exemplars^24–26^, and on parametrically varying single aspects of an object image at a time (e.g. by changing the size of the image^27–29^, or its position on the screen^25,28,29^). This contrasts with our experience in the real world, where each encounter we have with a specific object involves multiple simultaneous visual transformations caused by changes in factors such as lighting, viewing angle and distance, context, and object shape (for non-rigid objects).

**Figure 1.**
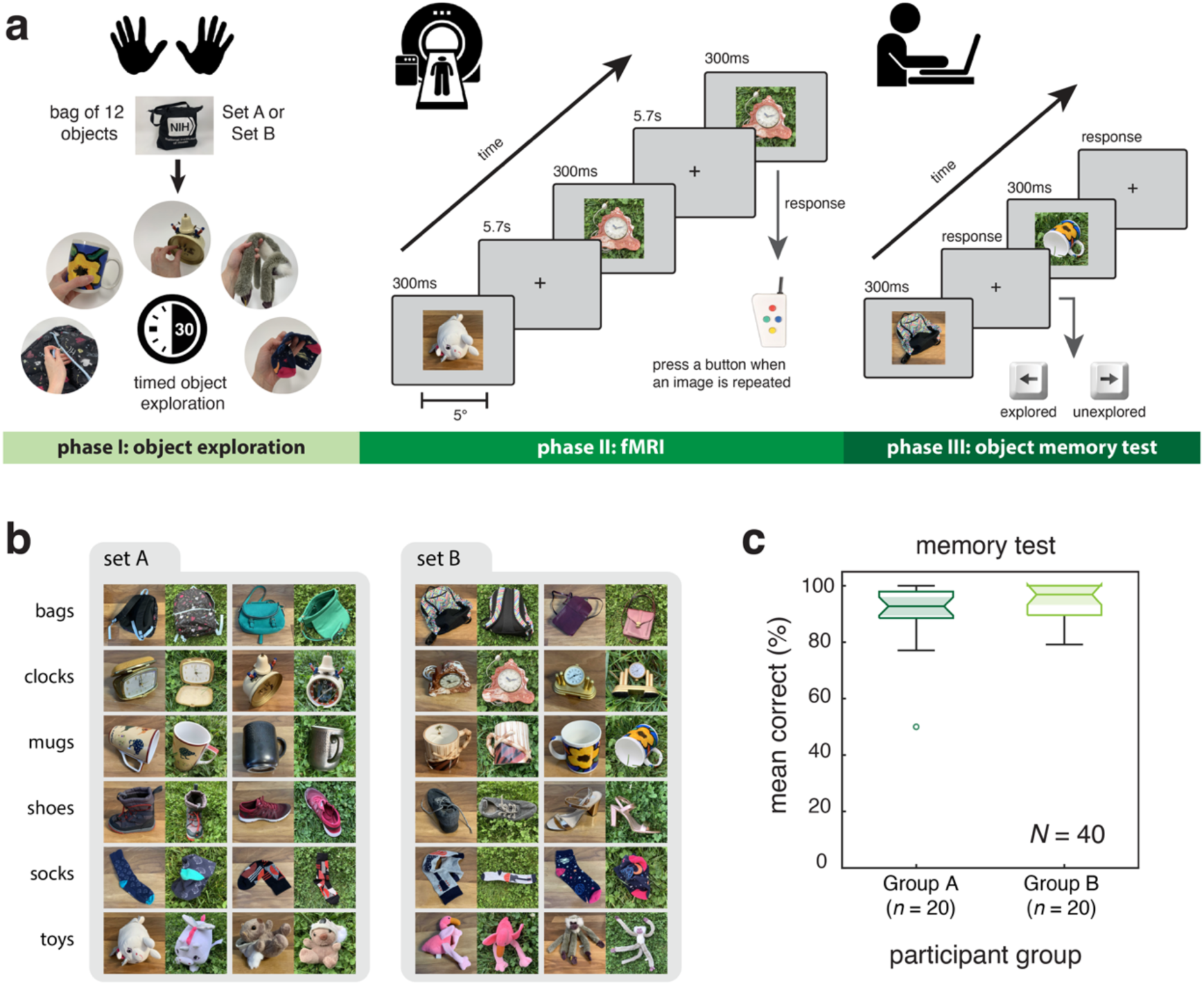
Experimental design and methods. **(a)** In phase 1 of the experiment, participants (*N* = 40) physically explored the 12 real-world objects from one of the object sets (Set A or Set B) one at a time for 30 seconds each. In phase 2, participants’ brain activity was recorded using event-related task fMRI while they looked at pictures of the 24 objects from both sets, half of which they had previously explored. Each object was depicted twice in different contexts for a total of 48 images as shown in (c). In phase 3, participants completed a behavioral memory test in which they viewed each of the 48 images used in the fMRI experiment and indicated whether they had explored the depicted object or not. **(b)** The 48 images used in the fMRI experiment. Each set had two unique exemplars from six object categories, for a total of 12 objects. Each object was photographed in two different contexts (indoor and outdoor) for a total of 24 photographs per object set. All participants saw all 48 images in the fMRI experiment, regardless of which object set they had explored. They did not see any of the photographs prior to the fMRI experiment. **(c)** Mean accuracy on the post-scan object memory test for all participants as a function of whether they explored Set A (Group A, n = 20) or Set B (Group B, n = 20) objects prior to the scanning session. Box plots show the median (center line), upper and lower quartiles (box edges) and the nonoutlier maximum and minimums values (whiskers) for each participant group. The shaded notches for Group A and Group B overlap, indicating no significant difference between the medians at the *p* < .05 level. Memory performance was near ceiling for both groups, indicating that they remembered which objects they had explored and recognized them from the photographs.

To preview the results, we found that object identity could be successfully decoded from object-responsive regions of cortex on the lateral occipital and ventral temporal surfaces even across different photographs of the same object— providing strong evidence for invariant object representations. Object representations in ventral occipitotemporal cortex were not modulated by real-world experience, but they were modulated by visual experience, with suppression of the BOLD response over repeated presentations of the same object image. Real-world exploration of an object was associated with a strong response in cortical areas outside the ventral visual pathway, including posterior cingulate and medial parietal cortex. Our results are important in constraining models of object recognition, and emphasize that in object-responsive cortex, object representations are highly visually-driven. In contrast, viewing a picture of a previously experienced object additionally engages medial brain regions associated with learning and memory— even across modalities and outside the context of an explicit memory or learning task. Together, the results show that the visual representation of an object and its associations engage different regions in the human brain, highlighting the importance of considering the role of cortical areas outside of visual cortex in visual object perception.

## Results

To investigate the effect of real-world experience with an object on the neural representation of its photographic image, we created two stimulus sets of everyday objects (bags, clocks, mugs, shoes, socks, toys). Each set of 12 objects had two unique exemplars from each of the six categories, which were photographed in two different contexts (indoor and outdoor) for a total of 24 images per set (**Fig. 1b**). Participants (*N* = 40) freely explored the 12 objects from one set (either Set A or Set B) one at a time for 30 seconds each prior to scanning (**Fig. 1a**). Following exploration, we recorded participants’ brain activity using fMRI while they viewed all 48 photographs of the 24 objects, half of which they had explored. Participants had not seen any of the images prior to the scan, yet a post-scan memory test revealed that they were able to recognize which objects they had explored from the photographs (**Fig. 1c**). There was no significant difference in the mean accuracy on the memory test for participants who explored Set A objects (*M* = 89.90%, *SD* = 11.77%) compared to those who explored Set B objects (*M* = 94.58%, *SD* = 5.91) as shown by a two-tailed independent samples t-test (*t*_(38)_ = −1.592, *p* = 0.120, 95% CI diff: [-10.649, 1.274], *d* = −0.503).

### Highly invariant object representations in object-responsive cortex

First, we established that explored and unexplored objects produced an equivalently strong magnitude of response in object-responsive regions within lateral occipital (LO) and ventral temporal cortex (VTC) (**Fig. 2a**). LO and VTC were functionally defined in each participant’s native brain space using data from independent localizer runs which included pictures of scrambled versus intact objects that were not used in the main experiment (**Fig. 3a**). A within-subjects 2×2 repeated measures ANOVA confirmed that there were no significant main effects of region of interest (*F*_(1,39)_ = 0.107, *p* = .745, *η*_p_^2^ = .003) or object exploration (*F*_(1,39)_ = 0.802, *p* = .376, *η*_p_^2^ = .020), and no significant interaction (*F*_(1,39)_ = 0.690, *p* = .411 *η*_p_^2^ = .017). Additionally, we were able to successfully decode the object images from the patterns of BOLD activation in areas LO and VTC for all object pairs (**Fig. 2b**), regardless of whether the depicted objects had both been explored or not (explored object pairs— LO: *t*_(1,39)_ = 9.578 *p* < .0001; VTC: *t*_(1,39)_ = 8.768 *p* < .0001, unexplored object pairs— LO: *t*_(1,39)_ = 8.620 *p* < .0001; VTC: *t*_(1,39)_ = 8.663 *p* < .0001, object pairs with one explored and one unexplored object— LO: *t*_(1,39)_ = 9.721 *p* < .001; VTC *t*_(1,39)_ = 9.694 *p* < .001; Bonferroni correction (*k* = 6) applied for multiple comparisons at the *α* = 0.05 level). Decoding performance was higher in magnitude overall for area LO than VTC (*F*_(1,39)_ = 22.664, *p* < .001, *η*_p_^2^ = .368) but there was no difference in decoding accuracy as a function of the type of object pair (*F*_(1.120, 43.684)_ = 0.096 , *p* = .786, *η*_p_^2^ = .002), and there was no interaction (*F*_(1.068, 41.669)_ = 0.466, *p* = .511, *η*_p_^2^ = .012). Mauchly’s test indicated that the sphericity assumption was violated for the main effect of pair type (*X*^2^*_(_*_2)_ = 58.506, p < .001) so the Greenhouse–Geisser correction was applied to these statistics and any interactions in the within-subjects 2×3 repeated measures ANOVA.

**Figure 2.**
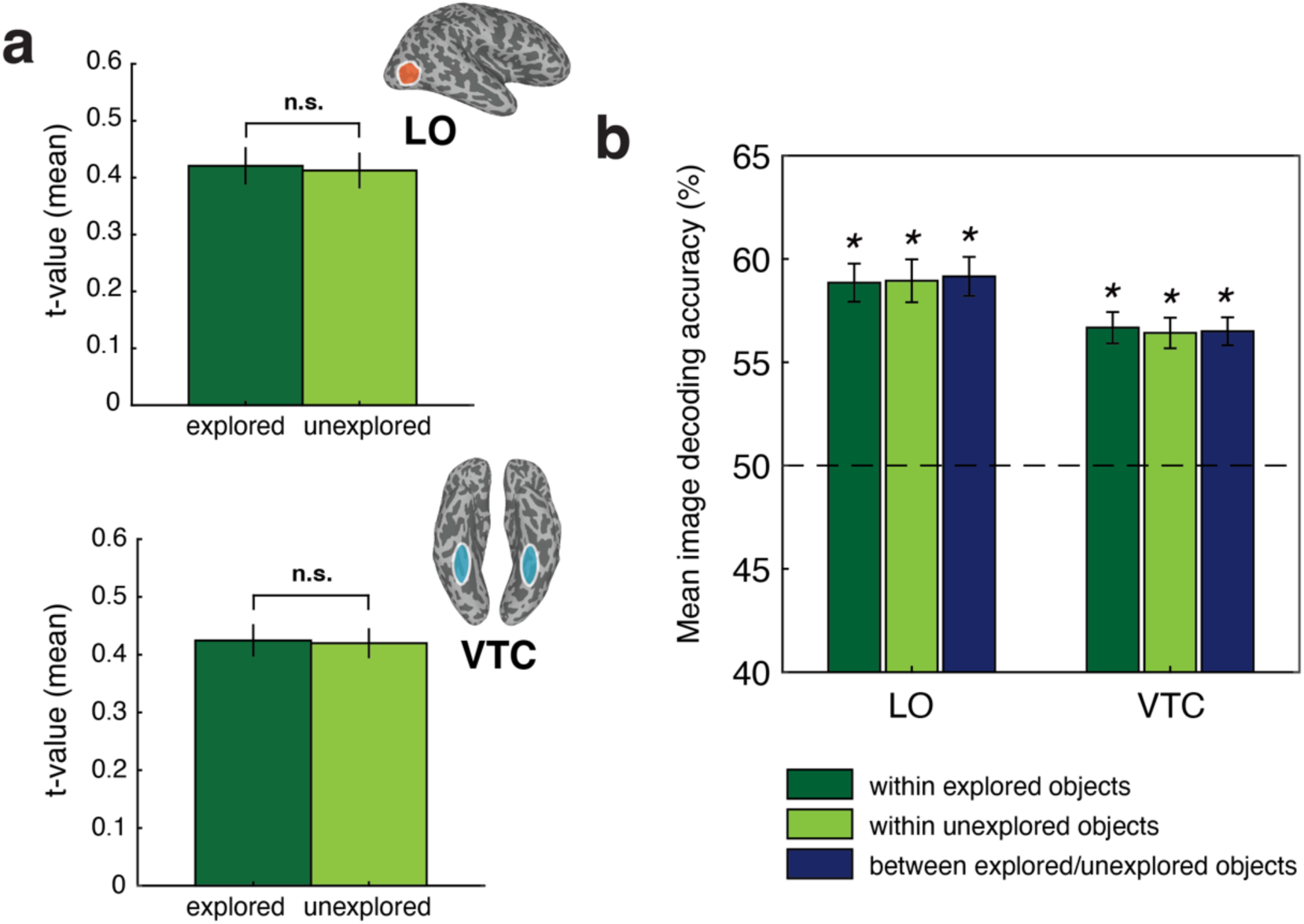
Response to explored and unexplored objects in visual object-selective areas. **(a)** The univariate response calculated as the mean t-value averaged across all voxels, object images, and participants (*N* = 40) is shown separately for object-responsive regions on the lateral occipital (LO) and ventral temporal (VTC) cortical surface. Insets show the location of the regions of interest, which were defined in each participant’s native brain space from independent localizer data. **(b)** Mean pairwise decoding of the object images across all participants (*N* = 40) and unique image pairs (*n* = 1,128) from patterns of BOLD activation in object-responsive regions LO and VTC as a function of whether both images in the pair were explored objects (*n* = 276 pairs), unexplored objects (*n* = 276 pairs), or one explored and one unexplored object (*n* = 576 pairs). The dashed line marks chance decoding accuracy (50%), asterisks indicate decoding performance significantly above chance (*p* < .0001, Bonferroni correction applied for multiple comparisons). Error bars are +/- 1 SEM.

**Figure 3.**
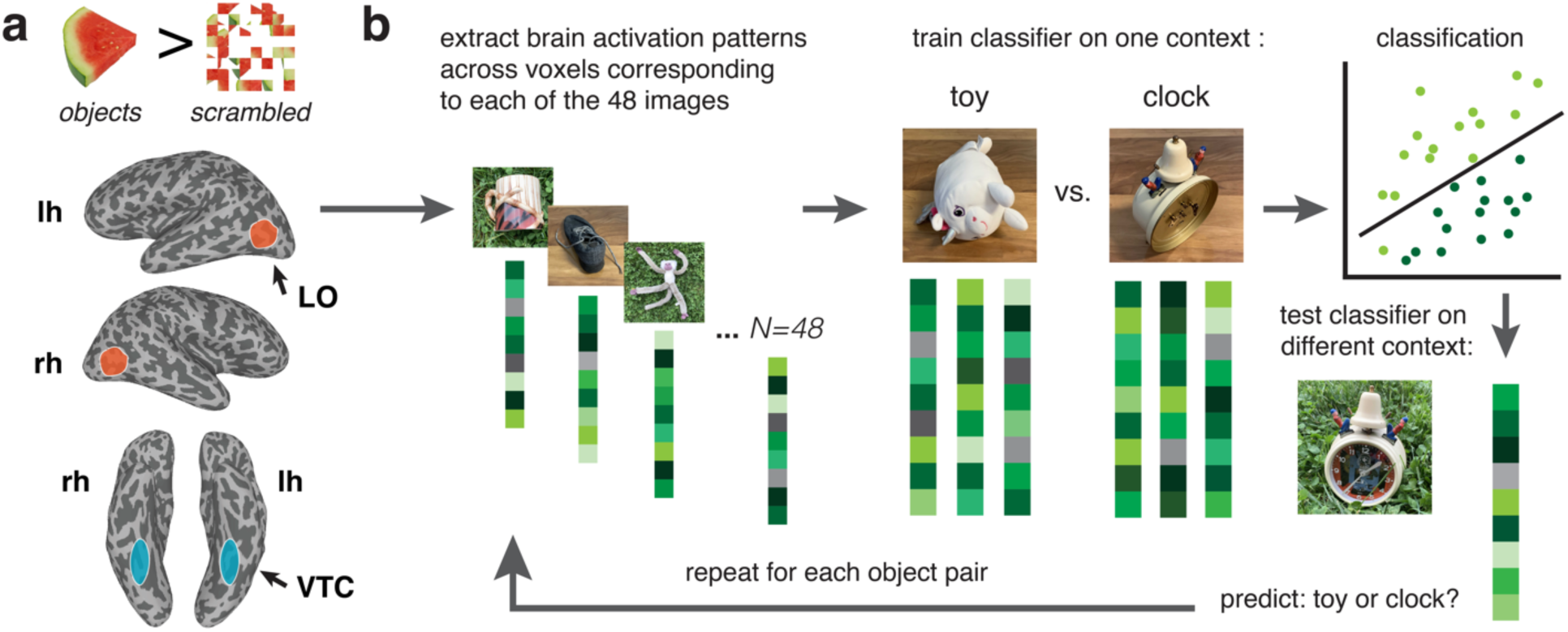
Method for cross decoding object identity across context from object-selective cortex. **(a)** Two regions of interest based on independent localizer data were defined in each participant’s native brain space on the inflated cortical surface from the contrast *objects > scrambled objects*. LO was defined as the cluster of voxels responding more to intact objects on the lateral occipital surface, and VTC was defined as the cluster of voxels responding more to intact objects in ventral temporal cortex. **(b)** Cross-decoding of object identity across context (indoor vs. outdoor scene) was performed by training a linear classifier to discriminate between a pair of objects photographed in one context (e.g. indoor) from the patterns of activation across voxels in a ROI and then testing the classifier on discrimination between the same pair of objects in the opposite context (e.g. outdoor). This procedure was repeated for each object pair.

Second, we found that object-responsive LO and VTC both contained highly invariant object representations of individual objects, even for those that were unexplored. To conduct a strict test for invariant object representations in object-responsive LO and VTC, we attempted to decode the identity of objects at the exemplar level across viewing context from patterns of brain activity measured with fMRI (**Fig. 3**). We used a cross-decoding approach and trained a linear classifier to discriminate between patterns of brain activity in these regions associated with looking at a pair of objects in one context (e.g. indoors), and tested classifier performance on new data from trials of the same object exemplars pictured in a different context (e.g. outdoors). Results of the decoding analysis are shown in **Fig. 4a** separately for each group of participants (Group A and Group B, defined by which set of objects the participants explored). One-tailed one-sample t-tests with the Bonferroni correction applied for multiple comparisons (*k* = 4) were used to assess whether decoding performance was greater than chance (50% accuracy) for each group. For both groups of participants, cross decoding of object identity across context was significantly above chance in object-responsive region LO for explored (Group A: *t_(19)_* = 6.536, *p* < 0.0001, 95% CI: [54.7516, Inf], *d* = 1.462 ; Group B: *t_(19)_* = 8.892, *p* < 0.0001, 95% CI: [55.6532, Inf] , *d* = 1.988) and unexplored objects (Group A: *t_(19)_* = 5.528, *p* < 0.0001, 95% CI: [54.8610, Inf], *d* = 1.236; Group B: *t_(19)_* = 5.462, *p* < 0.0001, 95% CI: [54.0800, Inf], *d* = 1.221). Cross decoding of object identity across context was also statistically significant for both participant groups in region VTC for explored (Group A: *t_(19)_* = 6.186, *p* < 0.0001, 95% CI: [54.1317, Inf], *d* = 1.403; Group B: *t_(19)_* = 6.009, *p* < 0.0001, 95% CI: [54.4920, Inf], *d* =1.344) and unexplored objects (Group A: *t_(19)_* = 9.252, *p* < 0.0001, 95% CI: [55.1826, Inf], *d* = 2.069; Group B: *t_(19)_* = 6.419, *p* < 0.0001, 95% CI: [54.6858, Inf], *d* = 1.435). These results indicate impressive generalization in LO and VTC— the identity of an object could be cross decoded across different photographs of the object, consistent with a high tolerance to visual feature differences in the underlying brain representations.

**Figure 4.**
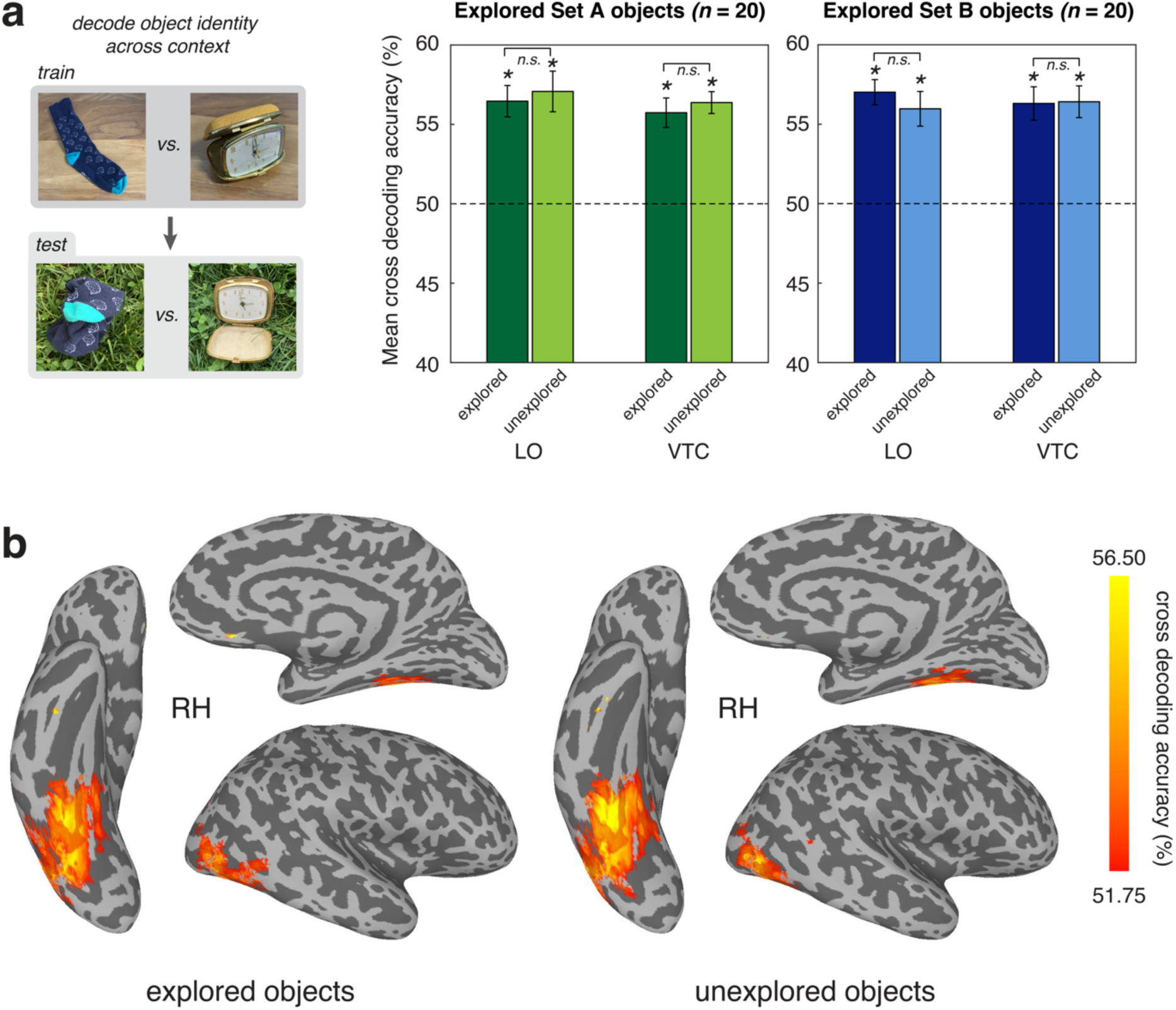
Cross decoding of object identity across context. **(a)** Results for cross-decoding object identity across context for object-responsive regions LO and VTC as a function of whether the object was explored in real life or not. Mean decoding accuracy is averaged across decoding iterations and participants. Error bars indicate +/- 1 SEM. Asterisks denote decoding performance above chance at the *p* < .05 level, Bonferroni corrected for multiple comparisons. **(b)** Whole-brain searchlight results for cross decoding object identity across context as a function of whether participants explored the object in real life. Mean decoding accuracy is averaged across decoding iterations and participants (*N* = 40). Only the right hemisphere is shown here for brevity, however the pattern of results was similar for the left hemisphere (see **Supplementary Fig. 1**).

Next, we tested whether there was a difference in cross decoding accuracy for objects that were explored versus those that were unexplored. For participants who explored Set A objects, there was no significant difference in the accuracy of cross decoding object identity across contexts between explored and unexplored objects in LO (*t_(19)_* = −0.635, *p* = 0.533, 95% CI: [-2.633, 1.407], *d* = −0.142, two-tailed) or in VTC (*t_(19)_* = −0.670 *p* = 0.511, 95% CI: [-2.635, 1.357] , *d* = −0.150, two-tailed). This result was replicated in both LO (*t_(19)_* = 1.335, *p* = 0.198, 95% CI: [-0.595, 2.691], *d* =0.298, two-tailed) and VTC (*t_(19)_* = −0.117, *p* = 0.908, 95% CI: [-2.018, 1.805] , *d* = −0.026, two-tailed) for participants who explored Set B objects. Bayesian analysis revealed moderate support for the null hypotheses that there was no difference in decoding performance for explored versus unexplored objects in LO or VTC for participants in both Group A (LO: *BF* = 4.833; VTC: *BF* = 4.730) and Group B (LO: *BF* = 2.575 VTC: *BF* = 5.824). Thus the invariant object representations in object-responsive cortex do not appear to be modulated by real-world exploration of the depicted objects.

To determine whether there were additional areas outside of our pre-selected object-responsive regions of interest that contained information about object identity that was invariant to context, we repeated the cross-decoding analysis as a whole-brain searchlight (**Fig. 4b; Supplementary Figure 1**). Consistent with our region of interest analysis, we found that cross-decoding of object identity across context was possible across large regions of lateral occipital and ventral temporal cortex in both hemispheres.

### Object exploration modulates activity in medial parietal cortex

In object-responsive visual cortex we found invariant object representations that were not modulated by real-world object exploration. To examine whether any cortical areas were modulated by prior real-world exploration of the depicted objects, we conducted a whole brain univariate analysis using a general linear model (GLM). An advantage of the counterbalanced experimental design is that the contrast *Explored objects > Unexplored objects* is defined as *Set A > Set B* for Group A participants (*n* = 20), and as *Set B > Set A* for Group B participants (*n* = 20). Thus when conducting a whole brain group level analysis on the entire sample (*N* = 40), any differences found cannot be due to specific images, and must be due to the differences in object exploration. From the group level analysis, we plotted the t-values for all voxels surviving cluster correction (see Methods) on the inflated cortical surface (**Fig. 5a**). This whole brain analysis revealed significant effects of object exploration in the posterior cingulate and medial parietal lobe in the left hemisphere (*t* = 3.55, *p <* .001). Similar effects were revealed to a lesser extent in the right hemisphere, with a significant cluster in the medial parietal lobe (*t* = 3.55, *p <* .001). The unthresholded maps on the cortical surface show similar patterns of medial activation in both hemispheres (**Fig. 5a**). We observe no significant clusters of differential activation for explored objects elsewhere, including somatosensory and motor regions. These data show that activity in medial parietal cortex in response to viewing a photo of an object is modulated by prior real-world experience with that depicted object.

**Figure 5.**
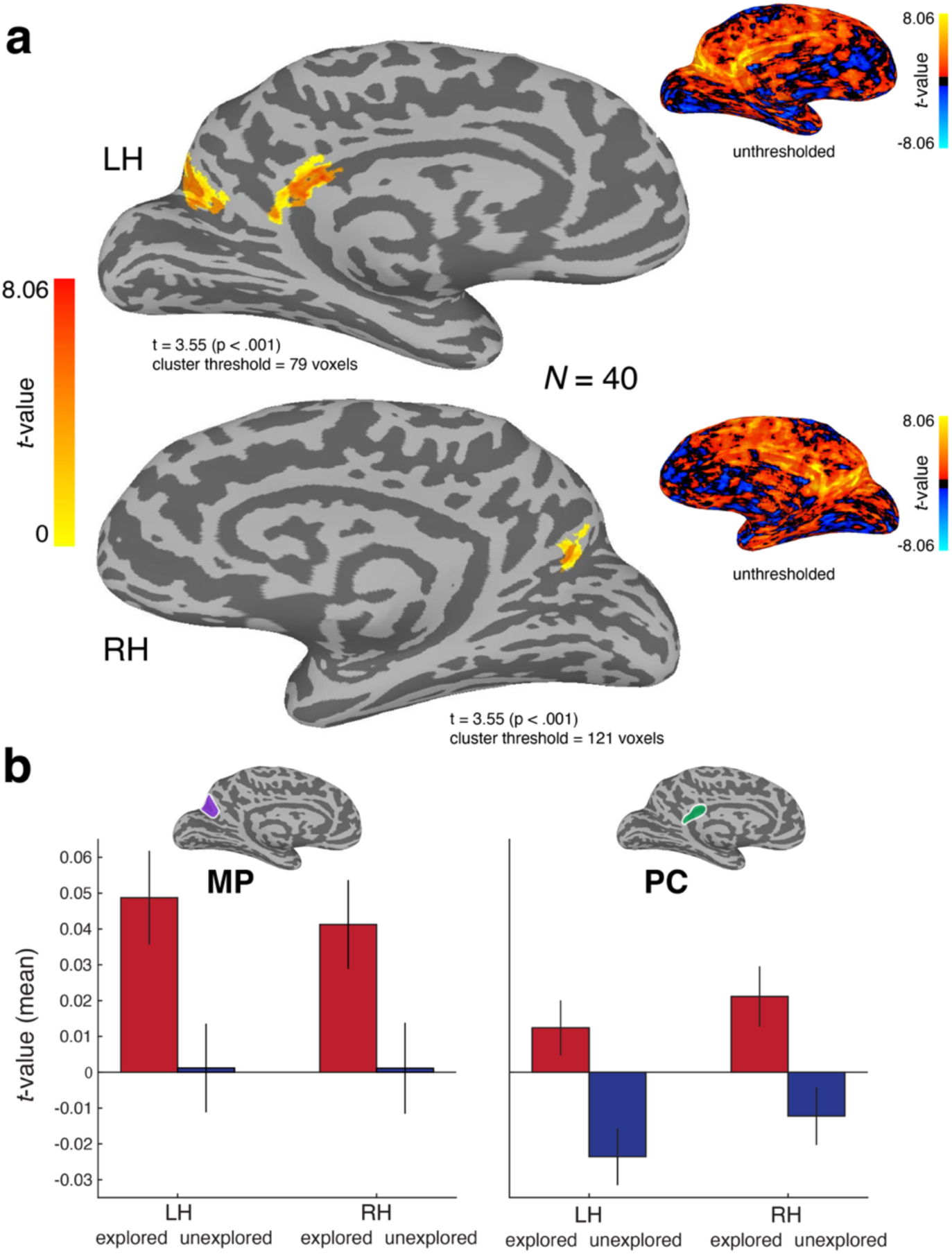
Whole brain response to viewing photographs of explored versus unexplored objects. **(a)** t-values from the GLM contrast (*explored - unexplored objects*) are plotted on the inflated cortical surface, the analysis was conducted on combined data for all *N* = 40 participants regardless of which object set they explored. For participants in Group A (*n* = 20) who explored Set A objects, this corresponds to the difference in activation for looking at pictures of *Set A - Set B* objects. For participants in Group B (*n* = 20) who explored Set B objects, this corresponds to the difference in activation for looking at pictures of *Set B - Set A* objects. As image set is counterbalanced across participants, any observed differences at the group level arise from the effects of object exploration. Only the medial view is shown for the left and right hemispheres as no other voxels survived cluster correction (*p* < .001). Insets show unthresholded group maps for each hemisphere on the medial surface. **(b)** Mean response to explored and unexplored objects as a function of hemisphere for medial parietal (MP) and posterior cingulate (PC) regions of interest. Insets show approximate location of MP and PC on the medial view of the inflated cortical surface. ROIs were functionally defined from the data for independent groups of subjects— see Methods for details.

We examined the response magnitude for explored and unexplored objects in these medial regions by defining two independent regions of interest in medial parietal (MP) and posterior cingulate (PC) cortex from the peaks of activation in one group of participants (e.g. Group A) and calculating the response magnitude from the data for the other group of participants (e.g. Group B— see Methods). We observed an increase in response for explored objects relative to unexplored objects bilaterally across both medial parietal and posterior cingulate cortex, with a stronger overall response in MP (**Fig. 5b**). This was confirmed with a within subjects 2×2×2 repeated measures ANOVA that revealed significant main effects of region of interest (*F*_(1,39)_ = 5.254, *p* = .027 *η*_p_^2^ = .119) and object exploration (*F*_(1,39)_ = 33.455, *p* < .001, *η*_p_^2^ = .462). Although there was no main effect of hemisphere (*F*_(1,39)_ = 0.641, *p* = .428, *η*_p_^2^ = .016), the interactions between hemisphere and region of interest (*F*_(1,39)_ = 4.512, *p* = .040, *η*_p_^2^ = .104) and hemisphere and object exploration (*F*_(1,39)_ = 4.549, *p* = .039, *η*_p_^2^ = .104) were both statistically significant, reflecting small differences in the patterns of response between hemispheres. No other interactions were significant.

### Repetition suppression in object-responsive cortex is not modulated by real-world exploration

We observed no modulation of activity in LO and VTC by real-world object exploration (**Fig. 2**), which suggests that exploring an object does not suppress the overall response to images of that same object. However, there may be a more nuanced change in response. The visual experience of handling the objects may lead to repetition suppression^30,31^, a reduction in response when viewing images of the objects. Such reduced responsiveness has been reported when images of an object are repeated^11,27,32^, although repetition suppression can be image-specific in visual cortex^33^ and may not generalize from 3D to 2D experience^20^. However, as repetition suppression is known to occur in object-responsive cortex for repeated presentations of the same stimulus^11,27,32^, it is possible that averaging the fMRI response over all presentations of the object images masked an earlier differential response for explored versus unexplored objects, particularly as participants had not seen any of the object images prior to the scan.

To determine whether such learning effects may account for the lack of modulation by real-world exploration, we plotted the mean response in LO and VTC to each object image over time, as a function of whether the object was explored or not (**Fig. 6a**). We observed classic repetition suppression for object images in both LO and VTC, but there was no modulation by real-world experience. We conducted a within-subjects 2 x 2 x 8 repeated measures ANOVA with the factors *ROI* (LO or VTC), *exploration* (explored vs. unexplored objects) and *run* (experimental runs 1-8) on the mean responses to each image across all voxels for the 38 participants who completed 8 experimental runs. The data from the remaining two participants who completed only 6 experimental runs (*n* = 1 from Group A, *n* = 1 from Group B) were excluded from this statistical analysis but are included in Fig. 6a. Mauchly’s test indicated that the sphericity assumption was violated for the main effect of run (*X*^2^ (27) = 60.083, *p* < .001) so the Greenhouse–Geisser correction was applied to the degrees of freedom for these statistics and any interactions. There was no main effect of ROI (*F*_(1, 37)_ = 0.209, *p* = .650, *η_p_*^2^ = .006) or object exploration (*F*_(1, 37)_ = 0.535, *p* = .469, *η_p_*^2^ = .014). In the visual ROIs, the response magnitude was similar for pictures of objects that had been explored compared to those that were not explored prior to the scan. Importantly, there was a main effect of run (*F*_(4.846, 179.310)_ = 2.916, *p* = .016, *η_p_*^2^ = .073), and a significant linear trend for run (*F*_(1,37)_ = 8.402, *p* = .006, *η_p_*^2^ = .185), indicating significant repetition suppression for both explored and unexplored objects in the object-responsive ROIs (**Fig. 6a**). The linear trend for the ROI x run interaction term was also significant (*F*_(4.846, 179.310)_ = 2.916, *p* = .016, *η_p_*^2^ = .073), indicative of the steeper slope for VTC. No other interactions were statistically significant. Thus in areas LO and VTC, the magnitude of response to a given image decreases as a function of repeated presentation across runs, regardless of whether that object was explored in real life or not.

**Figure 6.**
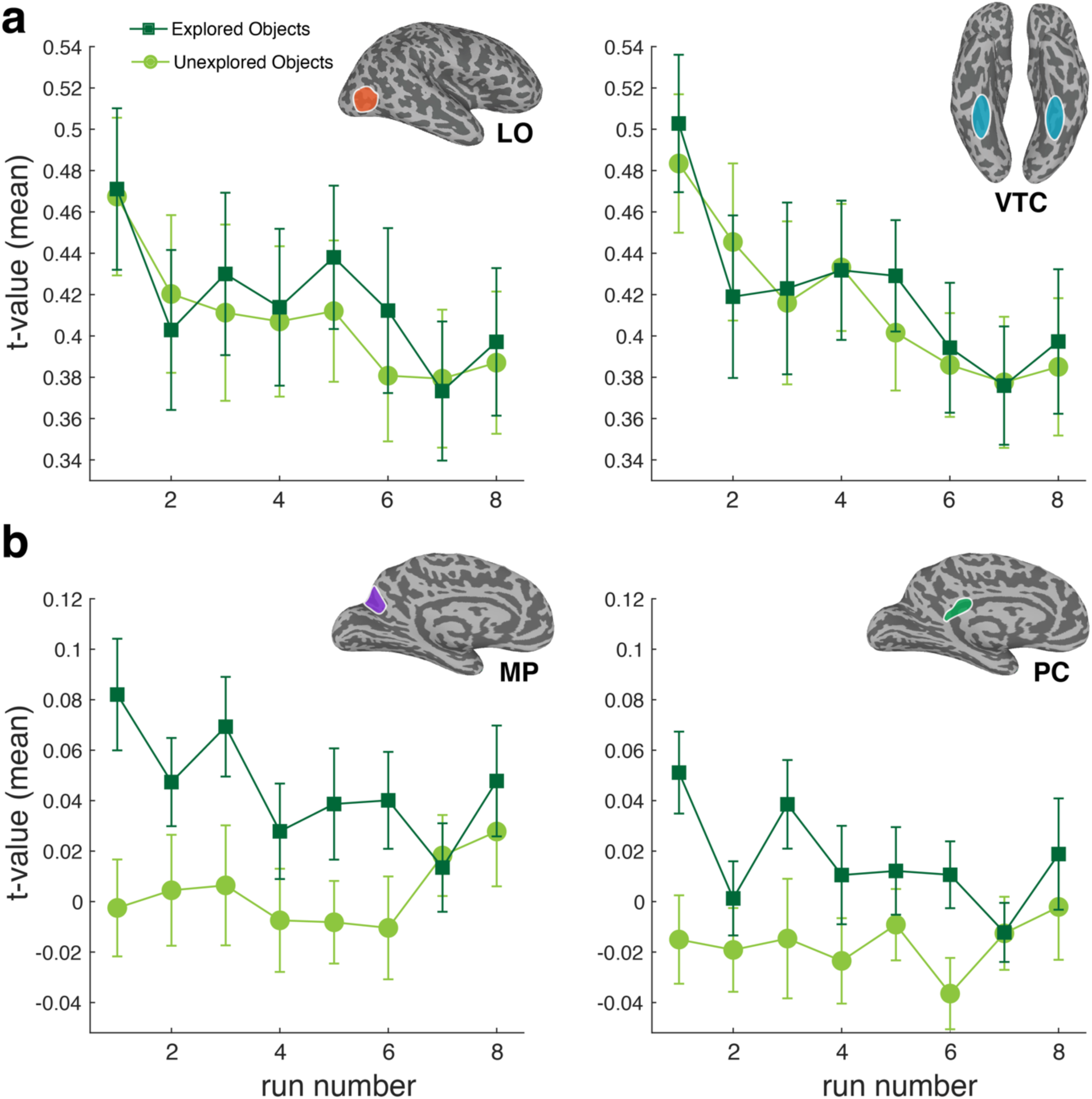
Differences in the response to the repeated presentation of object images over time in object-responsive and medial brain areas as a function of object exploration. The mean t-value averaged across all participants (N = 40) and the 48 images is shown as a function of run number (one presentation of each image per run) and plotted separately for images of explored (n = 24) and unexplored (n = 24) objects. **(a)** The object-responsive regions in the lateral occipital cortex (LO) and ventral temporal cortex (VTC) show repetition suppression for both explored and unexplored objects. **(b)** The medial parietal (MP) and posterior cingulate (PC) regions of interest show an enhanced response to explored objects compared to unexplored objects, which is evident from the first presentation of the images in the first run. Error bars show ± 1 between-subjects SEM.

### Medial parietal areas show enhanced responses to photos of explored objects

The medial parietal areas did show a differential response to explored objects, however we also checked for learning effects in these regions by plotting the mean response over time (**Fig. 6b**). Two new regions of interest — medial parietal (MP) and posterior cingulate (PC), were functionally defined bilaterally for each group of participants using independent fMRI data from the whole brain contrast for the other group (see Methods). As for the visual ROIs, we conducted a 2 x 2 x 8 repeated measures ANOVA with the factors *ROI* (MP or PC), *exploration* (explored vs. unexplored objects) and *run* (experimental runs 1-8). Mauchly’s test indicated that the sphericity assumption was violated for the main effect of run (*X*^2^ (27) = 48.418, p < .001) so the Greenhouse–Geisser correction was applied to these statistics and any interactions. There was a significant main effect of object exploration (*F*_(1, 37)_ = 31.311, *p* < .001, *η_p_*^2^ = .458), indicating an overall higher response magnitude for pictures of objects that had been explored compared to those that were unexplored. There was also a main effect of ROI (*F*_(1, 37)_ = 5.873, *p* =0.020, *η_p_*^2^ = .137), reflecting the stronger responses overall in MP (**Fig. 6b**). There was no main effect of run (*F*_(5.081, 187.995)_ = 0.967, *p* = 0.440, *η_p_*^2^ = .025), and none of the interactions between the main effects were statistically significant. There was some evidence of repetition suppression for explored objects but not for unexplored objects (**Fig. 6b**), supported by the absence of an overall linear trend for run (*F*_(1, 37)_ = 1.100, *p* < .301, *η_p_*^2^ = .029), but a significant linear trend for the interaction between object exploration and run (*F*_(1, 37)_ = 7.766, *p* = .008, *η_p_*^2^ = .173). Unlike the object-responsive regions LO and VO, the medial parietal regions exhibit relatively sustained response enhancement for explored objects relative to those that were unexplored. Finally, since there were two presentations of each object per run (i.e. the outdoor and indoor photograph of each object), we confirmed the pattern of results was similar as a function of object presentation number for both visual and medial ROIs (**Supplementary Fig. 2**).

## Discussion

Using fMRI, we found that prior real-world exploration of an object modulates activity in medial parietal regions in response to later viewing an image of the same object. Importantly, our counterbalanced experimental design ensured that these effects cannot be explained by differences among the images and objects and are attributable to an effect of real-world experience. Notably, real-world experience with an object did not modulate activity in object-responsive regions of lateral occipital and ventral temporal cortex. Our analyses confirmed that even on the first presentation of an individual photograph during the fMRI experiment, there was no difference in the magnitude of response to explored versus unexplored objects in LO or VTC. Instead, these regions were visually driven, containing highly invariant representations of individual objects as demonstrated by successful cross decoding across different photographs of the same object taken in different contexts (indoor vs. outdoor).

We discovered that viewing photographs of explored objects automatically engaged bilateral medial parietal and posterior cingulate cortex, even in the absence of an explicit memory task, and even though participants had never seen the photographs of the objects before. Similar cortical regions have been implicated in the “parietal memory network”^34^, which involves medial and lateral regions of parietal cortex associated with both memory encoding and retrieval. However, this is the first time to our knowledge that memory retrieval activity in the parietal memory network has been shown to generalize across modalities— in our paradigm, memory encoding was for real-world objects during the exploration phrase, while memory retrieval occurred spontaneously for photographs of the same objects viewed during the fMRI experiment. Participants had never seen any of the photographs prior to the fMRI scan, and the effect of exploration was evident on the first presentation. Additionally, the memory retrieval in our paradigm was spontaneous and implicit (i.e. recognizing a previously explored object from its photograph), rather than in the context of an explicit learning or memory task. Consistent with our finding of an enhanced response for explored objects in MP and PC, repetition enhancement (rather than suppression) is found in the parietal memory network ^34–36^. Our results suggest that medial brain regions are engaged by the associations created by prior real-world experience with an object, which are spontaneously activated by recognizing an explored object in a photograph. It is also notable that these effects were produced by only thirty seconds of free exploration with an object, evidence that these associations form rapidly and persist for at least a couple of hours following an object encounter. Our results suggest that modulation of activity in medial regions by object familiarity is neither stimulus nor task-specific, which implies that these regions may have a key role in real-world visual memory and recognition.

Our data show that medial parietal cortex responds more strongly to pictures of objects that we have real-world experience with than to pictures of unfamiliar objects, but there are several ways in which exploring a real object may alter the neural representation of its photographic image, and our data do not allow assessment of their relative importance. Exploration of a real-world object involves both tactile and visual sensory input, which may generate multisensory object representations^37,38^. In addition, exploration of an object generates an episodic memory of the experience, and participants’ thoughts during the exploration (for example, of what an object reminds them of, or their opinion of it) are another potential source of a differential response to explored versus unexplored objects. However, as we find no differential response in object-responsive visual areas, our data suggest that it is unlikely to be purely visually driven. Prior fMRI work has shown suppression of the response to familiar object images across object-responsive areas of ventral occipitotemporal cortex when participants were familiarized with a subset of images prior to the scan^39^. We observed classic repetition suppression in object-responsive regions to repeated presentation of the object photographs across runs, but no modulation by real-world experience. Thus repetition suppression in visual areas appears to be modality specific and depends on prior exposure to the same (or similar) image, rather than the same real-world object.

Perirhinal cortex has previously been associated with recognition memory and familiarity for object concepts and pictures^40–44^. In our paradigm, we did not observe a differential response in perirhinal cortex to pictures of explored versus unexplored objects, but there are several relevant differences between our paradigm and those that find perirhinal activation associated with familiarity. Many studies use concrete nouns (e.g. “apple”) as object concept stimuli^40,44^, in contrast to our stimuli which were photographs of specific object exemplars (e.g. a particular sock). They also frequently involve an explicit study phase followed by ratings of familiarity for the same stimuli^40,42–44^, whereas our participants completed a manual object exploration task and had never previously seen the photographs of the objects before the scanning session. Additionally, our paradigm did not involve an explicit memory or familiarity task. It remains an interesting question whether the familiarity signature in perirhinal cortex is modality-specific, but our data suggest that familiarity with a real-life object is not the same as familiarity for a specific image of that object.

An outstanding question is how the extent and nature of prior real-world experience with an object relates to the neural representation of its image. In our paradigm, the behavioral exploration of the objects depicted in the fMRI study was brief and controlled at thirty seconds per object. It is valuable to relate these results to prior studies that have used personally familiar visual stimuli, such as pictures of the participants’ own bag or office. The personal familiarity of places and objects has been linked to greater activation in medial regions including posterior cingulate cortex during a task in which participants judged whether an image was familiar or not^45^. Another fMRI study using images of personally familiar stimuli (e.g., images of a participant’s own vehicle or the face of their parent) found that their representation in inferior temporal cortex predicted the unique portion of an individual’s behavioral judgements of how similar the stimuli were^46^. Additionally, the results of an fMRI study involving novel objects created using the children’s construction toy K’NEX suggest that the way objects are used may alter their visual representation in the brain^47^. After using the novel objects to perform tasks (e.g. lifting blocks, crushing cups) in a training phase, activity increased in anterior parts of the left middle temporal gyrus, intraparietal sulcus, and premotor cortex during an in-scanner perceptual matching task in which participants judged whether two photographs depicted the same object or not. Participants were also faster at the behavioral task for the novel objects they trained with. Alongside our finding of strong activation in medial parietal cortex and the posterior cingulate for explored everyday objects, these studies highlight the importance of considering the effect of prior experience with objects on the neural representation of their image.

A complete understanding of human object recognition will require going beyond neural responses to images and detailing the neural representation of actual objects. Prior work has shown that real objects are processed differently than images of objects^5^. For example, repetition suppression in object-preferring regions of lateral occipital cortex and the posterior fusiform is weaker for real objects than for object images as measured with fMRI^20^. The neural representations supporting grasping movements for real objects also differ from those for photographs^19^. Behaviorally, real objects are more memorable than their photographs^15,16^ and capture attention more than images^18^, with infants as young as 7 months showing a preference for looking at real objects over pictures^17^. Patients with visual agnosia with impaired image recognition have also been shown to have preserved recognition for real world objects^48,49^. Our results complement these studies by showing that even very brief prior real-world experience with an object can modulate^48^ the neural representation of its image. Together these results demonstrate the value in incorporating more naturalistic behavioral paradigms and stimuli that go beyond the computer screen to understand human object recognition.

Our finding that object identity can be successfully cross decoded across very different photographs of the same object from patterns of activation across voxels in LO and VTC is important for establishing the invariance of these object-responsive cortical regions. Invariance is generally assumed to increase along the visual processing hierarchy^50^. Previous studies have shown some degree of invariance in object representations in human occipitotemporal cortex across singular visual transformations such as object size^27,28^, viewpoint^24^, and position in the visual field ^25,28^. Most earlier studies of invariance have focused on the category level rather than on the level of particular object exemplars^24–26,28^. We built on this work by conducting a strict test of object invariance for individual everyday objects. We were able to extract the representation of object identity from across different visual representations of each object by photographing actual objects in two different contexts (indoor and outdoor) and taking advantage of cross-decoding analysis methods to decode object identity across context. This test of invariance is much closer to how we experience objects in everyday life, with multiple visual features changing between the indoor and outdoor photographs of each object (e.g. luminance, contrast, viewpoint, distance, object shape [if non rigid], hue, illumination source, background, etc.). We found that both LO and VTC contained highly invariant representations of everyday household objects. Interestingly, we found no modulation of invariance by real-world object exploration in these visual areas. This may be because we used everyday objects such as mugs and shoes, and although participants had not encountered these specific exemplars before, they have a lifetime of experience with similar objects, which may be sufficient for supporting invariant visual representations of new exemplars. A study in monkey IT cortex reported a subset of neurons that responded to multiple views of novel objects that had previously been handled by the animal^51^. Although these responses were only able to be compared to unfamiliar novel objects for a total of four neurons, the data were consistent with more invariant representations for familiar than unfamiliar novel objects. This suggests that object novelty may be a relevant factor in the invariance of visual object representations.

In sum, our data show that the visual representation of an object is coded separately in the brain from the associations related to real-world exploration of that object. This is consistent with a framework that emphasizes that object concepts are not stored in a single brain region, but in networks of interconnected regions that extend beyond initial processing in sensory circuits^52^. Our discovery that ventral visual cortex contains highly invariant representations of individual objects that are not dependent on real-world experience underscores the efficiency of the human visual system in extracting object identity from the changeable visual features that accompany different viewing contexts. Real-world experience with an object has the potential to create a wide range of associations with an object from multimodal sensory details (how does it feel?) to context (where and when did you see the object?), associative memory (what does it remind you of?) and episodic memory (what did you do with the object?). Our data points to medial parietal cortex as an important associative region for the information gained from object exploration, complementing the visual object representations in LO and VTC. Overall, our study highlights the importance of considering the role of real-world experience in human object recognition. Although deep neural networks are now proficient in labeling objects from images^3,4^, human object perception extends beyond object labeling and encompasses a lifetime of experience manipulating and using objects to support our everyday behavior. A complete account of human object recognition will necessitate incorporating these human aspects of object perception into existing computational models.

## Methods

### Participants

The experiment was approved by the NIH Institutional Review Board (protocol 93-M-0170, clinical trials # NCT00001360). All participants gave written informed consent and received financial compensation for their time. Forty participants took part in the fMRI experiment (12 male, 28 female, age *M* = 24.68 years, *SD* = 4.80 years, range: 18 - 43 years). All participants were screened before taking part to ensure they were right-handed and neurologically healthy with normal or corrected-to-normal vision.

### Real-world objects

24 real-world objects were used as the experimental stimuli in the pre-scanning exploration task. They were divided into two sets of 12 objects, Set A and Set B. Participants explored only one of the object sets prior to scanning. Each set consisted of two exemplars from each of the following object categories: bags, shoes, mugs, clocks, socks, and stuffed toys. The object categories were chosen to be common household items that were small enough to be manipulable, and the specific object exemplars were selected to be distinctive enough to be recognizable in a photograph. The objects were sourced from the first author’s personal belongings, thrift stores, and the online marketplace Etsy.

### Visual stimuli

The visual stimuli were 48 high-resolution color photographs of the real-world objects. Each of the 24 objects was photographed twice in different contexts using an iPhone XS, once indoors (on a wooden floor) and once outdoors (on grass). The two photographs of each object were designed to be as visually different as possible, to separate out object identity from the visual features of the specific images as much as possible. The orientation of the object and its shape (if non-rigid, like a sock) were deliberately changed between the two photos. The original photographs were cropped square to equal size [600 x 600 pixels] with the object positioned in the center of the image. No other image manipulations were made. All participants saw all 48 object photographs in the fMRI experiment and post-scan memory test, regardless of whether they explored object Set A or Set B.

### Object exploration task

Prior to the MRI scan, participants completed a short object exploration task with one set of real-world objects (either Set A or Set B, counterbalanced across participants) and two experimenters. The first experimenter sat across a table from the participant and handed the participant each of the 12 objects one at a time from a black bag. Participants were given 30 seconds to explore the object however they chose. A second experimenter timed the exploration phase using a stopwatch, and the first experimenter returned the object to the bag at the end of the allocated time before producing the next one. The order of object presentation was randomized across participants, based on the order that the first experimenter pulled them out of the bag. The first experimenter recorded the object order on a sheet of paper by writing the corresponding presentation number [1-12] next to a picture of each object.

### fMRI data acquisition

The fMRI experiment consisted of a structural anatomical scan, two functional localizer runs, and up to eight experimental runs. MRI data were acquired with a 3.0T GE Discovery MR750 MRI scanner and 32-channel head coil at the Functional Magnetic Resonance Imaging Core Facility (FMRIF) at the NIH in Bethesda, MD. Functional scans were collected with a multiband three-echo EPI sequence with whole-brain coverage (TR = 2000ms, TEs = 12.9, 32.2, and 51.6 ms, flip angle = 75°, 90 x 90 acquisition matrix, voxel size = 2.5 x 2.5 x 2.5 mm, 50 slices). A high-resolution T1-weighted structural MRI scan (3D-MPRAGE sequence, 1 × 1 × 1 mm voxel size, in-plane matrix size: 256 × 256, 208 slices, TR = 6.952 ms, TE = 2.92 ms, FA = 8°) was also collected for each participant in the same session.

Visual stimuli were displayed using a flat-panel MRI-compatible 32′′ Cambridge Research Systems BOLDscreen with resolution 1920 × 1080 and viewing distance 2.5 m. Experimental and localizer stimuli subtended 5° × 5° of visual angle. Behavioral responses were collected using an MRI-compatible button box.

#### Object-localizer runs

Two independent functional localizer runs were used to define the object-responsive regions of interest in lateral occipital and ventral temporal cortex in each participant’s native brain space. The functional localizer stimuli were color photographs of objects and scrambled objects [600 x 600 pixels] from the BOSS database^53,54^. The scrambled object images were created using a custom-script written in MATLAB which defined an 8 x 8 grid over the original object image and shuffled the resulting 64 ‘tiles’ into a new location in the image. Each localizer run started and ended with a 16s fixation period, during which a black fixation cross was centered on a gray screen. The localizer runs used a block design, alternating between 16s blocks of objects and scrambled objects. There were 9 blocks of each stimulus type, for 18 blocks total. Within each 16s block, 20 stimuli were shown one at a time on a gray background in random order for 300 ms with a 500 ms inter-stimulus interval. Since the object photographs had a white background, a white square of the same size was shown on the gray background during the inter-stimulus interval to avoid unnecessary visual transients.

To maintain participants’ attention throughout the localizer runs, we used a standard 1-back task. Participants were instructed to press a key when they saw the same image presented twice in a row. In each block, 18 unique stimuli were shown, and 2 stimuli were randomly selected to be repeated for the 1-back task. There were two 1-back trials per block, and they were spaced so that one repeated image occurred at a random interval in the first half of the block, and the second repeated image occurred at a random interval in the second half of the block. Mean task accuracy across all participants and localizer runs was 86.56% (*SD* = 11.51%).

There was a total of 162 unique stimuli in each stimulus class (objects and scrambled objects), which were only shown once per run, except for the images repeated for the task trials. The images were randomly allocated to blocks each time the code was run. Each localizer run was 5.3 min long, and participants completed 2 runs during the MRI session — one before the experimental runs, and one after the experimental runs.

#### Experimental runs

In the experimental runs, we used an event-related design to measure the BOLD activation patterns evoked by each real-world object photograph. In each run, the 48 real-world object images (from both Set A and Set B) were shown once in a random order, with the restriction that no more than two objects from the same set were shown in a row. Each experimental run started and ended with a 16s fixation period — a black fixation cross was centered on a gray screen. Trials were 6 s in duration. On each trial, one object was shown for 300 ms on a gray background, and was then replaced by a black fixation cross during the 5.7 s inter-stimulus interval before the start of the next trial.

As for the localizer runs, we used a 1-back task to maintain participants’ attention throughout the experimental runs. There were four task trials per run, and four images were randomly selected to be repeated during the run (two from Set A objects, and two from Set B objects). Participants were instructed to press a key when they saw the same image presented twice in a row, and these repeated trials were removed from the analysis. Mean task accuracy across all participants and experimental runs was 88.75% (*SD* = 15.58%). Each experimental run was 5.7 min long, and participants completed 6-8 runs during the MRI session.

### Post-scan memory task

Following the MRI session, participants completed a short memory task outside the scanner on a MacPro laptop. This was to check that participants could remember and recognize the 12 objects they explored prior to the scanning session from the photographs used in the fMRI experiment. Each of the 48 photographs was shown one at a time on a gray background in the center of the screen for 300 ms. The order of images was random with the restrictions that (i) no more than two objects from the same set were shown in a row, and (ii) the two images of the same object (i.e. indoor and outdoor contexts) were not be shown in a row. Participants used the arrow keys to indicate whether they explored each object in real life (left arrow key) or not (right arrow key). The code waited for their response before proceeding to the next image after an 800 ms inter-stimulus interval. A black fixation cross was always displayed except for when an object image was on the screen.

### fMRI preprocessing

fMRI data were preprocessed using the AFNI^55^ software package. EPIs were slice-time corrected, motion-corrected, and co-registered to the participant’s individual anatomical volume using the customizable script *afni_proc.py*. Spatial smoothing of 4 mm full-width at half-maximum was applied to both localizer and functional runs. The multiecho EPI data was combined into a single time-series using AFNI’s “optimally combined” algorithm. All analyses were conducted in the native brain space of each participant.

### Functional ROI definition

Two functionally defined regions of interest (ROIs) were defined for each participant in their native brain space from the independent data in the functional localizer runs. Cortical reconstruction was performed using Freesurfer 6.0 from the structural scan for each participant. Inflated surfaces were visualized in SUMA^56^ for functional ROI definition, with the results of the GLM contrast (*objects > scrambled*) overlaid as t-maps. Object responsive regions were defined as the contiguous cluster of voxels that responded more to objects than scrambled objects in lateral occipital cortex (LO) and ventral temporal cortex (VTC) in each participant’s native brain space. Functional ROIs were defined on the inflated cortical surface for each participant and then transformed back into their native volume space for further analysis.

An additional two ROIs were defined from the response to explored versus unexplored objects in the experimental runs by splitting the participants (N = 40) into two groups (n = 20) based on which set of objects they explored (Group A and Group B) and defining the ROI for each group from the independent data from the other group. For each group, the unthresholded results of the GLM contrast (*explored > unexplored objects*) were overlaid as t-maps on an inflated surface in SUMA and the medial parietal (MP) and posterior cingulate (PC) ROIs were defined in each hemisphere as the region of contiguous voxels that responded more to explored objects in these anatomical locations. These ROIs were then transformed into the native volume space of each participant in the other group (i.e. Group A’s data was used to define Group B’s ROIs, and vice versa). The number of voxels in each hemisphere in each of the four ROIs is reported in Supplementary Table 1.

### fMRI multivariate pattern analysis

Decoding analysis was performed using The Decoding Toolbox (TDT)^57^ and MATLAB. Decoding was performed with a linear SVM in each ROI on the beta weights estimated in a GLM using AFNI for each of the 48 stimuli, producing a separate beta weight for each run. For the image-level decoding analysis (**Fig. 2b**), we used leave-one-run-out cross validation, and averaged accuracy over all cross-validation folds. For the cross-decoding analysis (**Fig. 4a**), the classifier was trained on object images in one context (e.g. indoor) and tested on object images in the second context (e.g. outdoor). We performed two-way classification for the cross-decoding (e.g. train on indoor, test on outdoor, and then the reverse) and averaged decoding accuracy over both versions of the analysis. For the whole-brain searchlight cross-decoding analysis (**Fig. 4b, Supplementary Figure 1**) we used the Newton linear SVM classifier implemented in TDT^57^ for efficiency, with a searchlight radius of 3 voxels. The searchlight was conducted in each participant’s native brain space, and then for visualization the results were mapped on to surface nodes in SUMA^56^ to produce the average group decoding maps for all participants (**Fig. 4b, Supplementary Figure 1**).

### fMRI group-level statistics

Statistical testing for the whole-brain univariate analysis examining the difference between explored and unexplored objects was performed using AFNI’s *3dttest++* and *SurfClust* functions on the standard mesh cortical surface. A group level t-test was performed for the contrast *Explored Objects > Unexplored Objects*. Note that for Group A participants (n = 20), this corresponds to *Set A > Set B*, and for Group B participants (n = 20), this corresponds to *Set B > Set A*. Cluster correction was applied to correct for multiple comparisons (at the *p* < .001 level) and the resulting thresholded t-statistics surviving correction for the whole sample (N = 40) were mapped on the inflated cortical surface (**Fig. 4**). The cluster threshold marking statistical significance at the *p* < .001 level was 79 voxels for the left hemisphere and 121 voxels for the right hemisphere.

## Notes

### Competing Interest Statement

The authors have declared no competing interest.

### Summary of Updates

new analyses added to Figure 2 and Figure 4, new Supplementary Figure added, revised main text to reflect new the analyses

