## Supplementary Information for "Brief encounters with real objects modulate medial parietal but not occipitotemporal cortex"

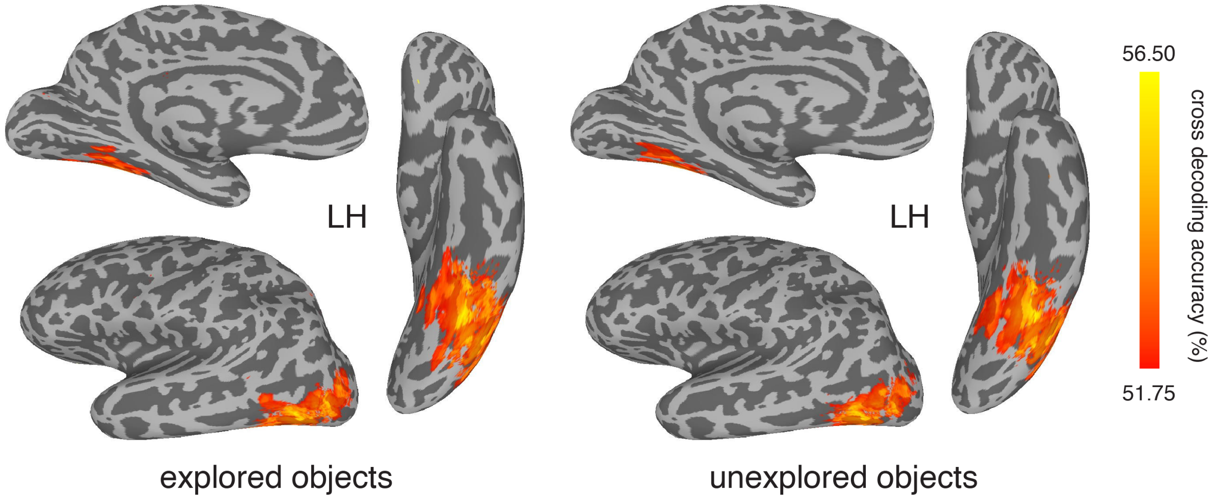


**Supplementary Figure 1.** Whole-brain searchlight results for cross decoding object identity across context as a function of whether participants explored the object in real life. As in Figure 4b, but shown here for the left hemisphere.

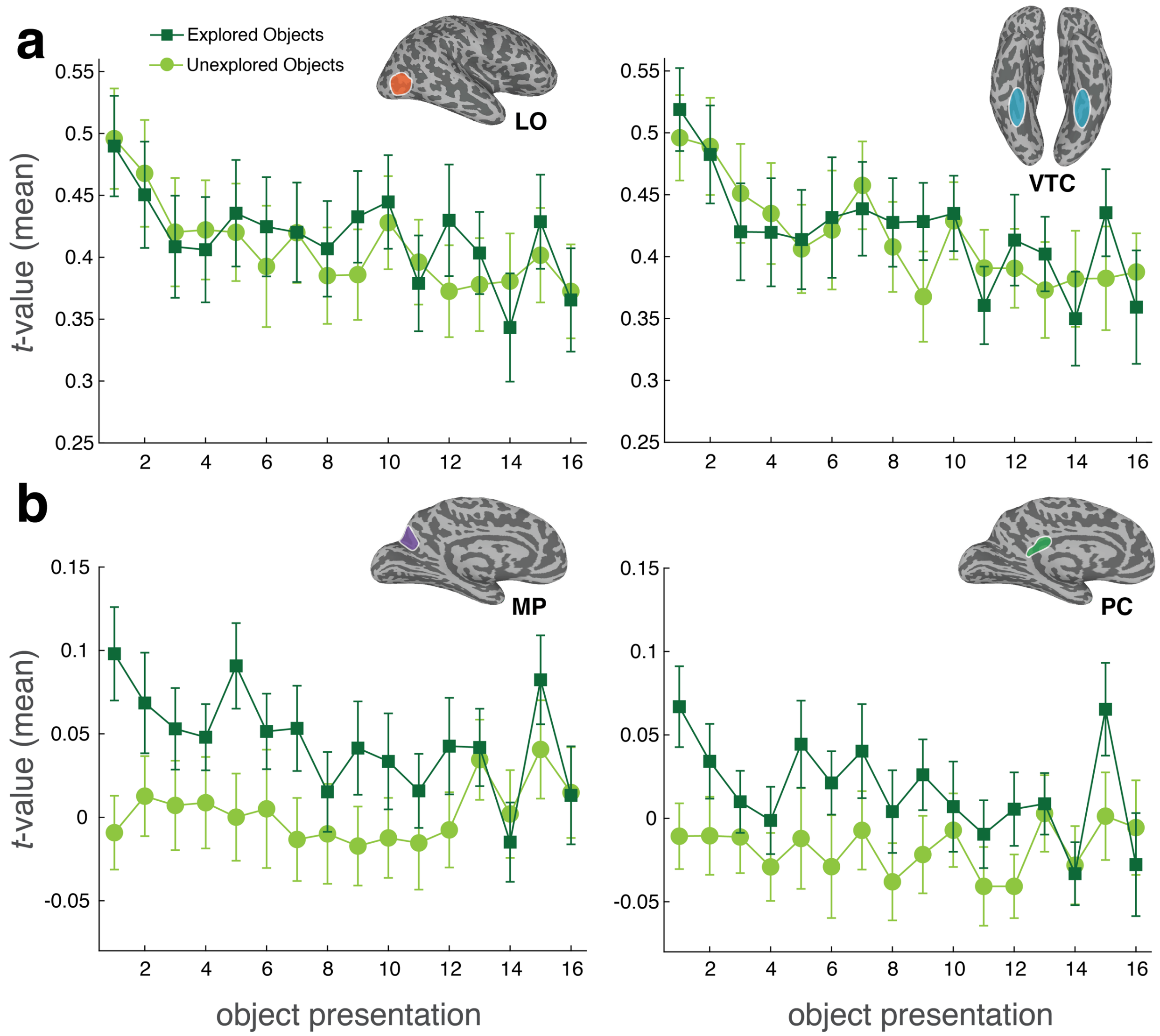


**Supplementary Figure 2. Differences in the response to the repeated presentation of objects over time in object-responsive and medial brain areas as a function of object exploration.** The mean t-value averaged across all participants (N = 40) and the 24 objects is shown as a function of object presentation number. Two presentations of each object occur per run, as there are two different images of each object. The data are plotted separately for images of explored (n = 24) and unexplored (n = 24) objects. **(a)** The object-responsive regions in the lateral occipital cortex (LO) and ventral temporal cortex (VTC) show repetition suppression for both explored and unexplored objects. **(b)** The medial parietal (MP) and posterior cingulate (PC) regions of interest show an enhanced response to explored objects compared to unexplored objects, which is evident from the first presentation of the object. Error bars show ± 1 between-subjects SEM.

**Supplementary Table 1. Mean number of voxels in each hemisphere for each region of interest, averaged across participants (*N*=40).**

| ROI | number of voxels rh | number of voxels lh |
|---|---|---|
| **LO** | 502 [*SD* = 122] | 528 [*SD* = 194] |
| **VTC** | 291 [*SD* = 105] | 306 [*SD* = 108] |
| **MP** | 506 [*SD* = 231] | 517 [*SD* = 266] |
| **PC** | 182 [*SD* = 23] | 205 [*SD* = 34] |
